# Multi-scale and multi-context interpretable mapping of cell states across heterogenous spatial samples

**DOI:** 10.1101/2024.08.31.610638

**Authors:** Patrick C.N. Martin, Wenqi Wang, Hyobin Kim, Henrietta Holze, Kyoung Jae Won

## Abstract

In clinical applications, spatial data collected under varying conditions, time points, or patients often lack discernible structural alignment. Computational tools designed to align adjacent tissue sections are unsuited for dealing with this structural heterogeneity. There is a growing demand for methods that can effectively align and compare spatial data in the absence of obvious visual correspondence. To address this challenge, we developed an interpretable cell mapping strategy by considering spatial context at various scales. Our approach outperforms existing mapping tools in dealing with heterogeneous samples and is flexible enough to map cells across samples, technologies, resolutions, developmental and regenerative time. Using our approach, we showed spatiotemporal decoupling of cells during development. We even performed alignment for a population of spatial data from cancer patients to identify sub- populations. Our interpretable mapping approach facilitates systemic comparison and analysis of heterogeneous spatial data.

## Introduction

From the initial discovery of cells through early microscopes in the 17^th^ century, we have come a long way in our understanding of cellular organization^1^. Today, spatial transcriptomics (ST) has become an increasingly popular means of probing cellular organization in relation to cellular identity^2–6^. ST measures the gene expression profiles as well as the location of cells within a tissue. For clarity, we will refer to spatial spots, barcodes, or indices simply as cells. Taken together, ST methods - in their varying flavors – can highlight spatially resolved gene expression patterns^7–13^. They can demonstrate how tumors interact with their microenvironment and how disease can be understood from a spatial perspective^14–17^. They can provide insights into cellular communication through ligand-receptor dynamics as well as cell-to-cell contact triggered expression modulations^18–21^. Importantly, ST emphasized how cellular identity should be viewed through the lens of spatial context^21,22^.

With the increasing availability and breadth of ST technologies, it comes to no surprise that they have become part of the biomedical research arsenal. As it is still difficult to obtain high quality spatial data and it is still expensive to produce, there are needs to compare the spatial data deposited in the public domain. However, data sets that are already available in a biological condition of interest might have been produced using different methodologies or technologies. In a clinical setting where a researcher might be interested in following the evolution of cells and their interactions across disease progression or as a response to treatment, it is difficult to expect tissues to share similar structures. We have even seen the emergence of 3D spatial stacks^9,23,24^ taken from adjacent tissues. A plethora of tools allowing the alignment of these tissue slices including PASTE^26^ and GPSA^28^ assuming the tissue structure similarity. But irregular nature of tumors across patients and conditions restricts their use in various circumstances. As such, the question remains: how do we compare the spatial context of individual cells when tissue structures do not match across samples or where sequential sampling is simply impossible?

Mapping single cell data to a spatial assay based on the transcriptome may suggest an alternative solution to align two or more ST assays based on their transcriptome^29,30^. For instance, Tangram uses a deep-learning approach to match sc/snRNA-seq to their estimated spatial location by maximizing the spatial correlation between scRNA and spatial data^29^. CytoSpace solves a linear assignment problem to match single cells to spatial locations^30^. In conjunction with single cell labels, CytoSpace can also map to high-resolution spatial data. However, since these methods were designed to map scRNA-seq to spatial data, the spatial context of cells is not or only partially considered. Mapping without spatial and biological context can lose the information related with tissue or cell microenvironment integrity. Algorithms such as Spatial-linked alignment tool (SLAT) provide a solution of spatial mapping by adopting a graph adversarial matching strategy to map cells between spatial samples with heterogeneity^31^. While SLAT considers the spatial context of cells across samples, it has not been tested systematically for tissues whose tissue structure is dissimilar. Also, its graphical representation does not provide clear interpretation of mapping factors. Moreover, all these mapping approaches cannot be applied to systemic comparison of hundreds of samples.

To address the challenge of mapping cells between structurally heterogenous tissues, we developed a novel approach to align two or even more spatial samples when they cannot be aligned well using classical approaches. For this, we performed mapping while considering spatial context in in the framework of a linear assignment problem. Specifically, we designed the cost matrix to minimize the pair-wise summation of multi- scale and multi-context spatial matrices including cell similarity, niche similarity, and spatial territory similarity (see Methods). Explicit use of context specific matrices increases the interpretability of cell mapping results and provides flexibility for mapping across technology and samples collected over time. We highlight the performance of our approach using synthetic data sets in a variety of scenarios with increasing complexity. We demonstrate how our flexible framework can effectively map cells between data sets, even between samples generated from different technologies, and spatial resolutions. We map cells across developmental and regenerative time in high-resolution Stereo-seq data. This allowed us to highlight genes involved in brain development and regeneration which were only present in a limited subset of cells. Additionally, the cost for cellular mapping provides a metric to cluster samples at the population level. The clustering results demonstrated how cellular context can recapitulate broad clinical metrics. Our cell mapping strategy grants us the ability to investigate biological differences between samples with unprecedented flexibility and interpretability.

## Results

### Mapping cells and their context across samples

With the increased availability of ST data sets, we aim to provide a flexible framework to compare spatial data by mapping a query data set onto a reference data set across conditions, technologies, and sections (Figure-1a). To achieve this, we formulate the matching of cell pairs as a Linear Assignment Problem (LAP) where the goal is to match “a worker” to “a tasks” which minimizes the overall cost (Figure 1b). To solve the assignment problem we used the Jonker-Volgenant algorithm^32^ In this instance, a “worker” is a cell from the query data whereas the “task” is a cell from the reference data (see Methods). The overall cost of mapping cells is defined by a pair-wise summation of score matrices across biological features (Figure 1c). We account for transcriptional similarities between cells, their niches, their spatial tissue territory, cell type labels, and the cell type composition of their niche. In addition, we offer a straightforward method to add custom and context specific matrices. The total cost matrix can be constructed from any combination of these biological features. For instance, while cell type labels can be used, they are not strictly required to map cells between samples. Our algorithm has been included in the Vesalius R package – a tool designed to retrieve spatial territories by applying image processing methodologies to spatial omics data^22^ (Figure- 1d). The territory similarity matrix is computed using Vesalius’s tissue territory detection capabilities. The new and improved functionalities of Vesalius provide a framework for spatial mapping of samples, mapping of cells across time and across technologies, and to cluster hundreds of spatial samples.

**Figure 1:**
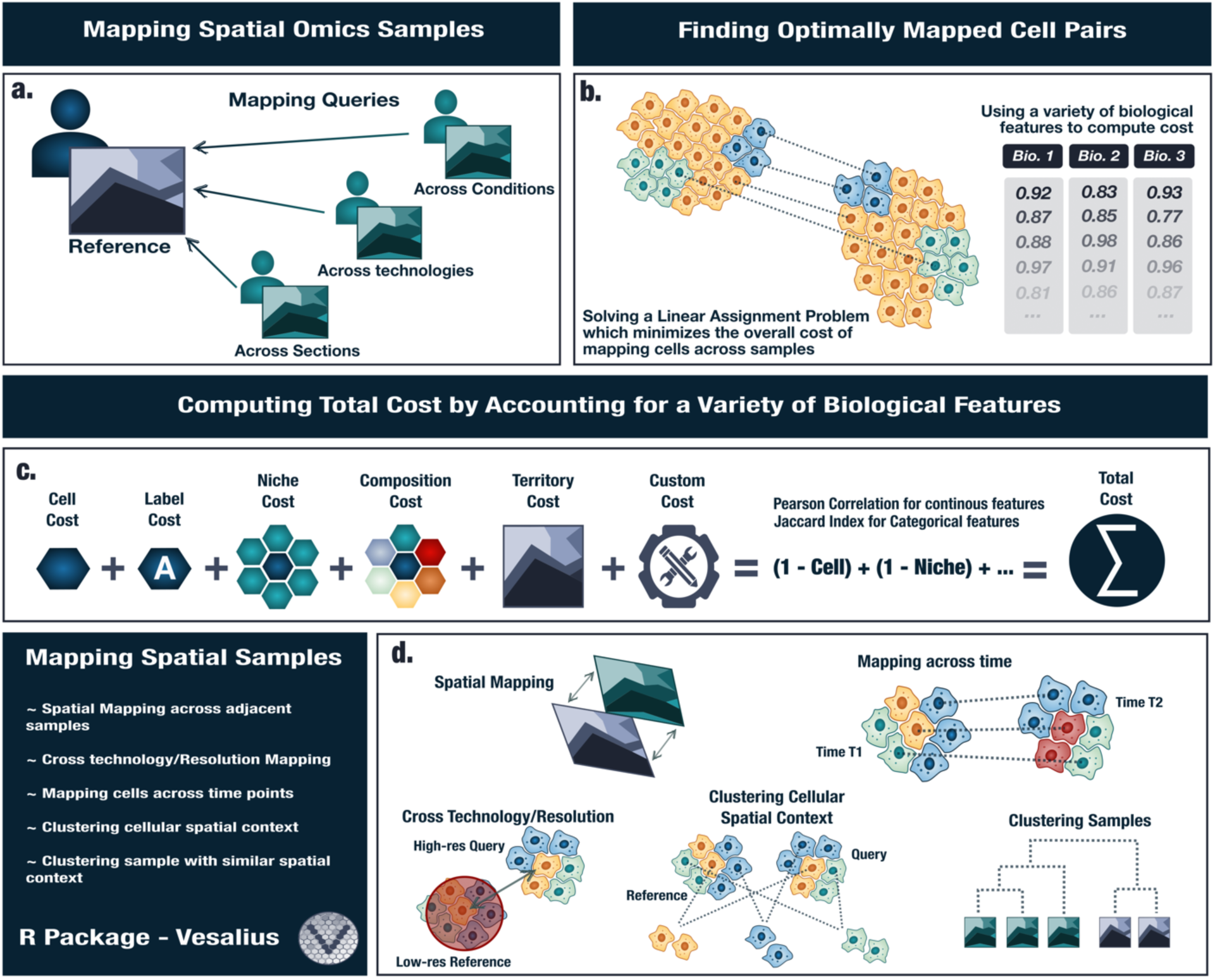
Overview of Vesalius’s cell state mapping strategy. (a) Sample across conditions, technologies, and sections are likely to present a heterogenous structure (b) Vesalius maps cells across samples by solving LAP while minimizing the mapping cost. To account for spatial context during mapping, Vesalius leverages a multitude of biological features. (c) The total cost matrix is a pair-wise summation of reciprocal similarity scores including cell similarity, label similarity, niche similarity, composition similarity, territory similarity, and even custom similarity scores. (d) Vesalius maps cells across samples, across time and, across technologies. It also performs clustering population of patient spatial samples.

### Benchmarking mapping performance in synthetic spatial data

To demonstrate the effectiveness of our mapping strategy, we elected to use synthetic spatial data since this will provide a robust ground truth (See methods). We used 3 spatial regimes (*circle*, *layer,* and *dropped*) that mimic the complexity of biological scenarios (Figure 2). In the *layer* regime, we aim to reproduce the cell state variability one might observe in a tumor in which expression patterns differ from the tumor micro- environment interface and its center. The *dropped* regime contains these layers with the added complexity that some cells may not be shared between data sets.

**Figure 2:**
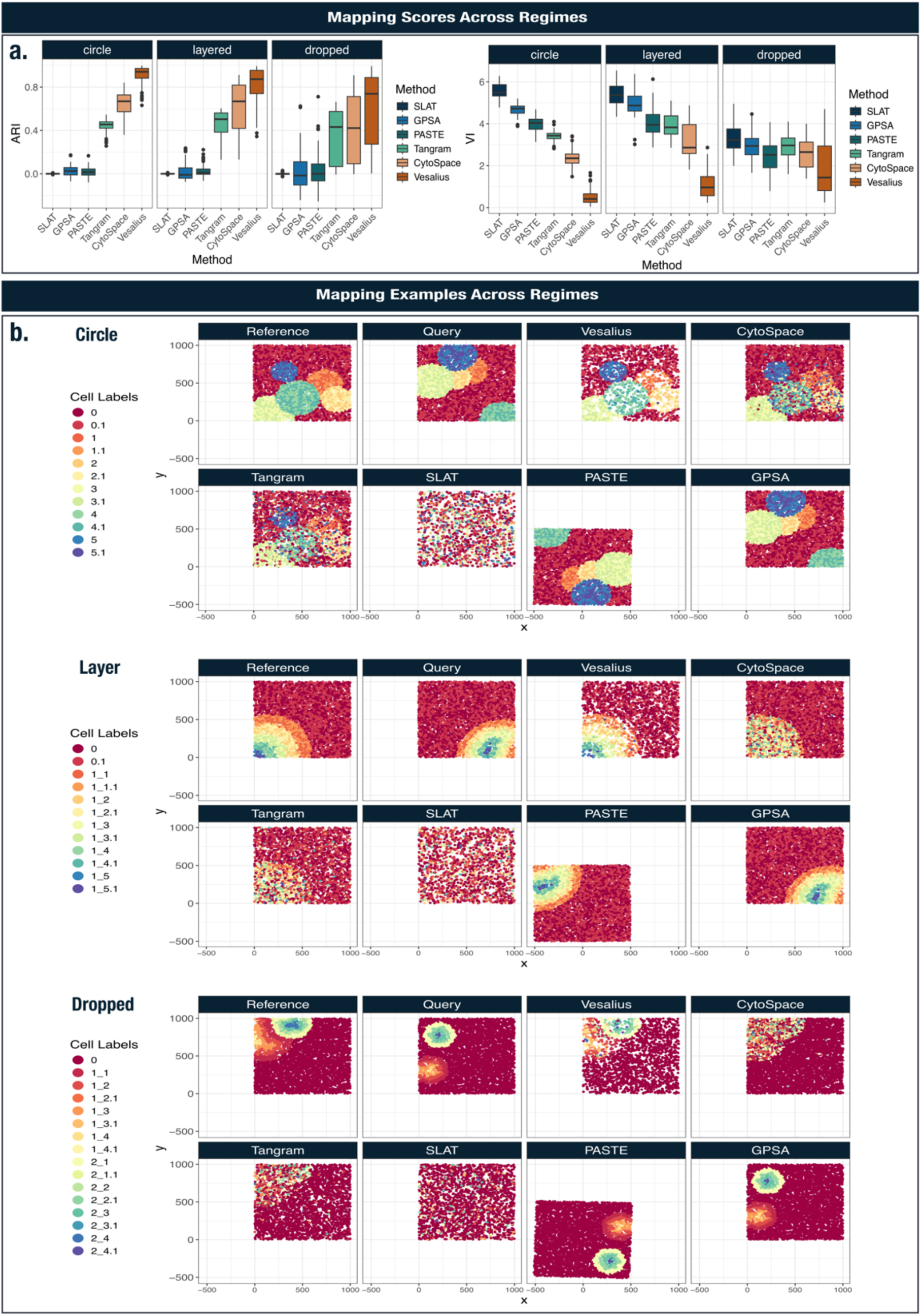
Benchmarking mapping performance in synthetic spatial data. (a) Using the cell type labels between samples, we computed the Adjusted Rand Index (ARI) and Variation of Information (VI) to quantitatively assess mapping performance. Vesalius outperforms other tools in mapping cells between heterogenous samples. (b) Example of mapping of the query cells onto the reference cells. Vesalius accurately mapped most cells across samples while method such as CytoSpace and Tangram -lacking spatial context – struggle to recover the cell’s spatial distribution. PASTE and GPSA only tried to rotate and distort the sample. SLAT was unable to accurately map cells between samples.

With these simulated data sets in hand, we compared the mapping performance of Vesalius using two scRNA-to-spatial mapping tools (CytoSpace^30^ and Tangram^30^) and spatial mapping tools including SLAT^31^, PASTE^26^, and GPSA^28^. For benchmarking, we constructed the total cost matrix from cell similarity, niche similarity, and territory similarity (obtained with Vesalius territory detection).

Overall, Vesalius performed competitively against other methods across metrics and simulation regimes (Figure 2a). The slight decrease in performance in the *dropped* regime could be attributed to the changes in the cell type composition of the neighborhood. CytoSpace and Tangram mapped cells between query and reference in the *circle* regime with reasonable accuracy but struggled to map cells in the *layered* and *dropped* regime where the expression difference between layers is more spatially subtle. PASTE and GPSA only rotated or distorted the query and were unable to handle samples structurally different each other (Figure 2b). SLAT performed poorly and was unable to accurately map cells in any regime even though it was designed to map heterogeneous spatial data

### Context specific mapping allows for accurate cell mapping across biological samples

We first mapped cells between similar tissue samples in Slide-seqV2^8^ mouse hippocampus data and mouse embryo seqFISH data^33^. In seqFISH murine embryo data, We elected to construct the total cost matrix from cell similarity, niche similarity, and niche composition as the transcriptome and the cell types are well annotated already. The type of the mapped cells by Vesalius matched well with the corresponding cell type in the reference including the forebrain/midbrain/hindbrain regions and cardiomycytes (Figure 3a). Since the total cost is constructed from individual biological features, we further investigated the contributing factors for mapping performance by investigating individual scores. The cell neighborhood composition score (Jaccard Index) shows the highest correspondence which tells that many cells in the mapped results share strong similarity in their neighborhoods composition (Figure 3b). It is also noticeable that the cell and niche similarity scores contribute to discriminating spatial context beyond cell type labels (Figure 3b – Figure S1c). We investigated how different cost matrix combinations modulate the mapping performance in seqFISH (Figure S1c) and found that the additional information provided by each cost matrix improves mapping performance.

**Figure 3:**
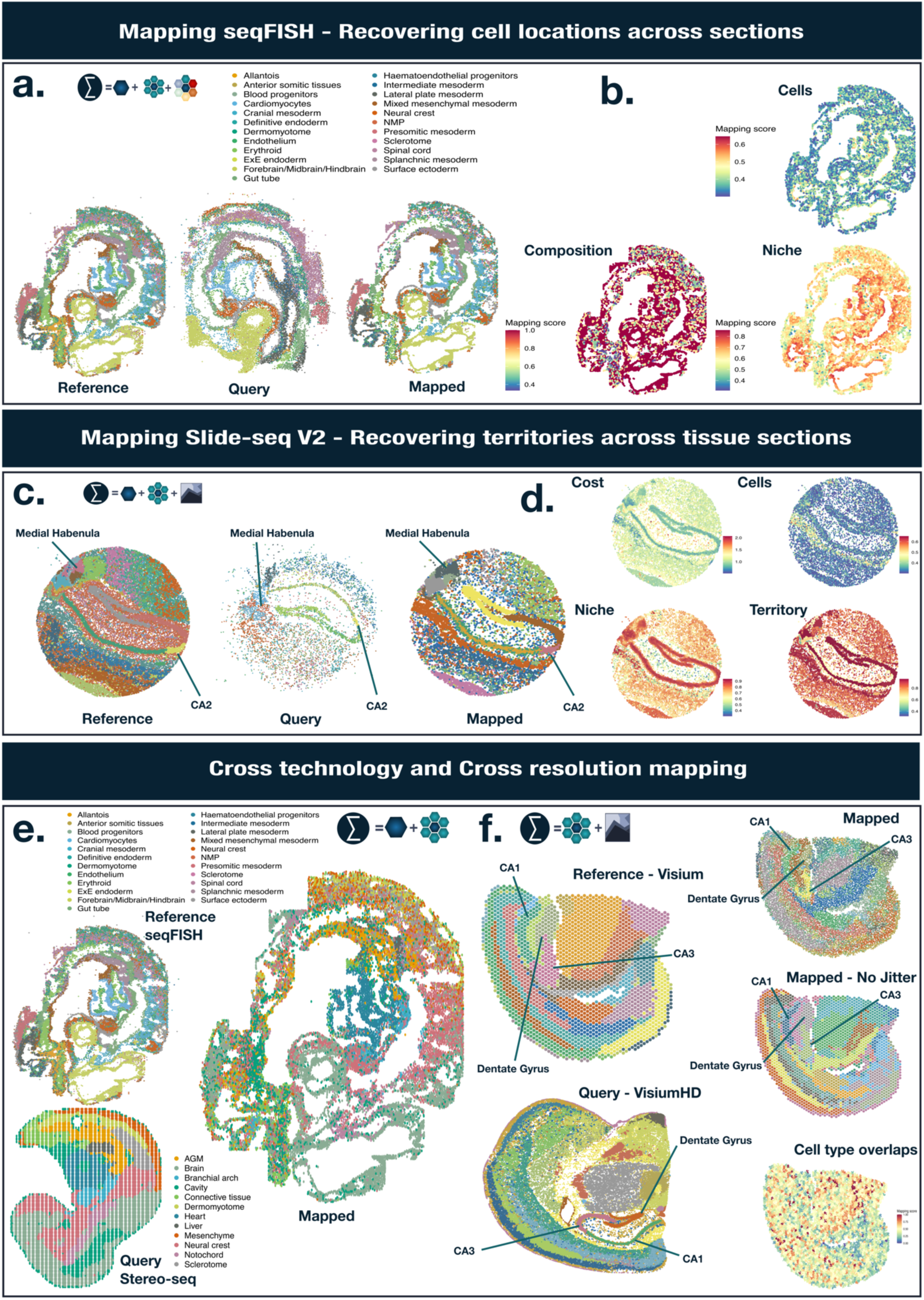
Cell Mapping across samples, technologies, and resolutions. (a) Mapping of seqFISH mouse embryo cells across samples (b) composition score, niche and cell similarity contributed to the mapping cost. (c) Mapping of cells in Slide-seqV2 taken from 2 separate mouse hippocampus sections. The tissue regions such as 2 medial habenula compartments and the CA2 field are recovered in the reference, the query, and in the mapped query cells. (d) The mapping cost and the scores for each context. The niche and territory similarities exhibited better discriminative value than cell similarity. (e) Mapping Stereo-seq mouse embryo cells onto seqFISH mouse embryo cells. (f) Mapping VisiumHD mouse brain onto Visium mouse brain. The tissue territories detected by Vesalius in the reference, the query, as well as the mapped cells with a jitter added to the coordinates. The cell type overlap using Jaccard index between estimated cell types in Visium spots and the VisiumHD.

In Slide-seq V2, we constructed the cost matrix from cell similarity, niche similarity, and territory similarity after removing low quality beads as that information is readily accessible for this data (see methods). We noticed that Vesalius recovers many tissue territories in both samples including the CA2 field and medial habenula compartments (Figure 3a).

After mapping query cells onto the reference, we called tissue territories and accurately recovered the expected tissue structures (Figure 3c). It is intriguing to see that the medial habenula is clearly found in the mapped results even though it is unclear in the query sample. We visualized the mapping scores associated with mapped cells (Figure 3d). While lower cost indicates better mapping, high mapping scores (Pearson’s correlation) indicates better correspondence between query and reference cells. Niche and territory scores showed high correlation and highlighted how these metrics aided in spatial mapping.

### Mapping cells across technologies and resolutions

More often than not, data sets of interest are generated using different technologies and yet we would still like to be able to mine insights from them. We tackled this challenge using our flexible mapping strategy. First, we mapped high-resolution Stereo-seq mouse embryo^9^ (E9.5) onto image-based seqFISH^33^ in the same tissue at a slightly earlier developmental stage (E8.75). While both data sets contain cell type labels, these differ greatly between data sets as is often the case when using data from different sources. As such we forwent the cell type labels, and we constructed the total cost from cell similarity and niche similarity only. Overall, the broad cell type labels matched well for the mapping across technologies (Figure 3e). For instance, brain and heart cells in the Stereo-seq are mapped to Forebrain/Midbrain/Hindbrain and Cardiomyocytes in seqFISH, respectively. We show the cell mapping results for Stereo-seq to all 3 seqFISH sections in Supplementary Figure S2. We also performed the converse mapping (seqFISH mapped to Stereo-seq) (Supplementary Figure S3).

To further test mapping across spatial resolution, we mapped VisiumHD mouse brain onto Visium of the same tissue (See data availability). We elected to use the 8 μm bins since this would strike a balance between high-resolution and reduced noise at each spatial location. We detected tissue territories in both samples (Figure 3f) and recovered expected tissue structures such as the CA1 field, CA3 field, and the Dentate Gyrus. To account for the difference in resolution between samples, the total cost matrix was constructed using niche similarity and territory similarity. More specifically, for each cell, we define the niche by using a distance radius around the center cell (see Methods). We defined the radius as the radius of a Visium spot to ensure that a Visium niche would only contain a single spot while visiumHD would contain all spots within that radius. Since mapping of high-resolution onto low-resolution data signifies that multiple spots will be mapped to the same low-resolution spots, we added random noise (jitter) to the coordinate values to better visualize the mapping results. Alternatively, we omitted the jitter to create pseudo-Visium data where each mapped cell will be merged into a mini-bulk count data at the spot level (Figure 3f).

We discovered that we were recovering the expected tissue structures, demonstrating that we can effectively map cells across resolutions. Interestingly, we observed finer tissue territories in the mapped region onto Visium assay compared with the reference.

To further investigate the cell types matching well between the two technology, we used RCTD^34^ to estimate cell type proportions in both Visium and VisiumHD using the same reference scRNA-seq data set^35^. We computed a Jaccard index between cell types associated to Visium spots and the mapped cell identities from VisiumHD. Overall, our results show a reasonable correspondence in mapped cell types between Visium and VisiumHD (Figure 3f). We do not observe distinct spatial patterns in the label overlap score indicating that there is no bias towards mapping specific cell types.

### Mapping cells across time in brain regeneration and embryo development highlights spatio-temporal expression patterns among mapped cells

The STOmics database provides a wide variety of high-resolution Stereo-seq data collections. Most notably are the MOSTA^9^ and ARTISTA^24^ data collections which provide spatial-temporal maps of murine embryo development and Axolotl brain regeneration. Using these collections, we applied our approach to mapping cells across developmental time. First, we focused our attention on brain regeneration with the aim of mapping cells forward in time. We selected 20 Days Post Injury (DPI) as the reference onto which we would map 15 DPI (Figure 4a) with a cost matrix constructed from cell similarity, niche similarity, and territory similarity. We observed that cells are accurately mapped in the 20DPI brain (right side of the spatial assay). Ependymoglial cells (ECG) – equivalent to neural stems cells – form a mass of reactive ECG cells (reaECG) where regenerative intermediate progenitor cells (rIPC3 and rIPC4 cells) will form at a later stage (Highlighted in figure 4a by blue circle).

**Figure 4:**
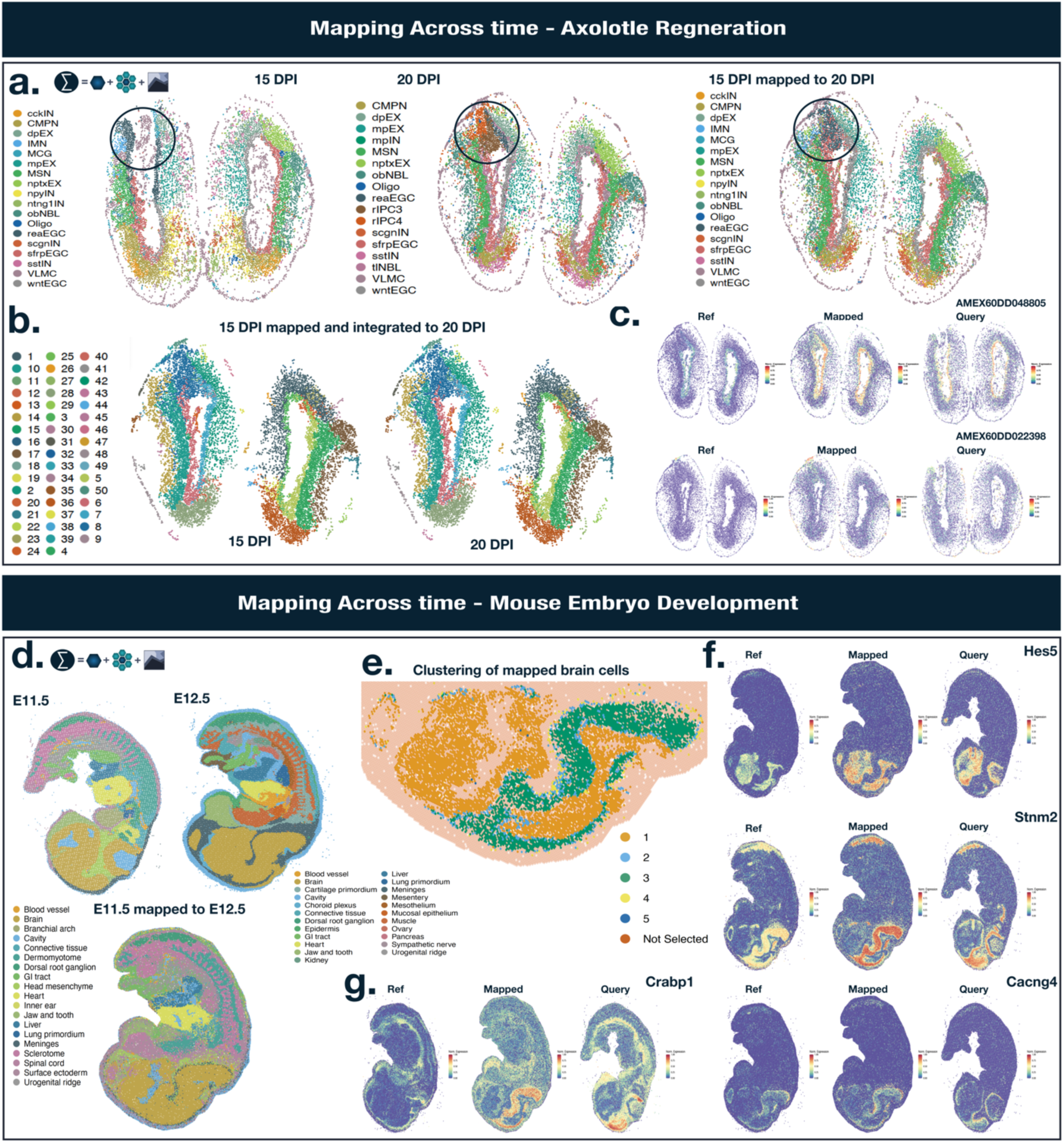
Cell Mapping across time. (a) Mapping Stereo-seq Axolotl brain regeneration data sets (ARTISTA) at 15 DPI to 20DPI. Blue circle highlights ependymoglial cells and their transition to regenerative intermediate progenitor cells. (b) Integrated tissue territories of mapped cells and reference cells. (c) Two differentially expressed genes at the zone of injury (territory 8 – dark blue): AMEX60DD048805 and AMEX60DD022398. (d) Mapping Stereo-seq Mouse embryo (MOSTA) development at stage E11.5 to stage E12.5. (e) Clustering of query brain cells. Brain cells are demarcated into sub-territories including cells that lay at the interface between brain regions (cluster 2) and the interface between the brain and other tissues (clusters 4). (f) Differential gene expression between clusters exemplified by *Hes5*, *Stnm2*, and *Cancng4* (g) *Crabp1* – a gene involved in the regulation of stem cell differentiation – shows a spatial-temporal decoupling behavior.

We created an integrated image stack (see Methods) upon which we can apply image processing and isolate common territories between reference and query data sets (Figure 4b). The left assay represents the mapped cells while the right represents the reference cells. The temporal alignment provides a unique opportunity to identify differentially expressed genes focusing on associated cell pairs. We computed differentially expressed genes between reference and mapped cells at the injury zone (territory 8 – dark blue) to uncover which genes were involved in the regeneration process. Out of the 123 differentially expressed genes, we found 2 notable genes AMEX60DD048805 (EDNRB) and AMEX60DD022398 (ARPP19). The former – homologous to the EDNRB human gene involved in vasculature and cell proliferation- is expressed in a subset of vascular leptomeningeal cells (VLMC) in the outer layers of the brain (Figure 4c). The latter – a gene involved in the cell cycle is expressed in a specialized set ECG cell, srfpECG and wtnEGC. The full list of differentially expressed genes is available in Supplementary Table 1.

We applied a similar process of mapping forward in time in mouse embryo. As an example, we selected developmental stage E12.5 as the reference data and E11.5 as the query data. The total cost matrix was constructed using cell similarity, niche similarity, and territory similarity. Once again, our approach shows a remarkable ability to map cells across space and time (Figure 4d). We aimed to explore how cell populations preferentially mapped to the same set of cells in the reference data. To achieve this, we performed hierarchal clustering (see Methods) using cell similarity using the brain cell in the query data and identified 5 clusters (Figure 4e). We observed the spatial distributions of the mapped cells: Cluster 2 and 4 highlight that cells were mapped at the interface of these brain regions (Figure 4e). We obtained the genes enriched for each cluster. For instance, *Hes5*, *Stnm2*, and *Cacng4* were differentially expressed in cluster 1,3, and 4 respectively (Supplementary Table 2). Spatial gene expression patterns for each cluster indicate that our mapping strategy recovers expected gene expression patterns across space and time (Figure 4f). Intriguingly, we discovered examples of spatiotemporal decoupling (Supplementary Table 3). For instance, *Crabp1*(Cellular Retinoic Acid Binding Protein 1), a gene involved in stem cell proliferation and differentiation^36,37^, shows high expression in cluster 3 of the query cells but is not expressed in the reference data.

### Sample stratification through cellular context mapping in proteomics data

We further applied our strategy to align hundreds of spatial samples. We randomly sampled 100 patients from Breast Cancer Image Mass Cytometry (IMC) data containing a total of 37 protein markers^38^. We conducted an initial filtering step to ensure that all data sets contained clinical level metrics such ER Status (Positive or Negative), ERBB2 positive, cancer grade, death, cancer subtype (PAM50) and years to outcome. We then selected samples which contained between at least 1000 cells. From the remaining data sets, we used ER Status to selected patient sub-populations since this metric enables balance and interpretability to our patient sampling procedure. We then mapped each sample to all other samples (excluding self-mapping) using a cost matrix generated from cell similarities, niche similarities, territory similarities, cell label, and nice composition. From each mapping event, we extracted the average cost and mapping scores to define overall sample mapping performance.

We selected best mapping pairs based on lowest overall cost (Figure 5a). Across all pairs, cell similarity, niche similarity, and territory similarity exhibited high average mapping scores. Niche composition had the lowest overall score. A large fraction of the best matching pairs also showed high correspondence with clinical metrics with 80 % having at least 3 out 5 metrics correctly matched (Figure 5b). ERBB2_pos was the most well predicted feature for our cost function followed by ERStatus. The least well predicted was cancer sub-type (PAM50) with 49 out 100 correctly predicted. However, this metric contains 5 sub-categories (Basal, HER2, Luminal A, Luminal B, Normal-like) making our predictions well above a random baseline. The majority of samples show a low difference in years to outcome between query and reference samples.

**Figure 5:**
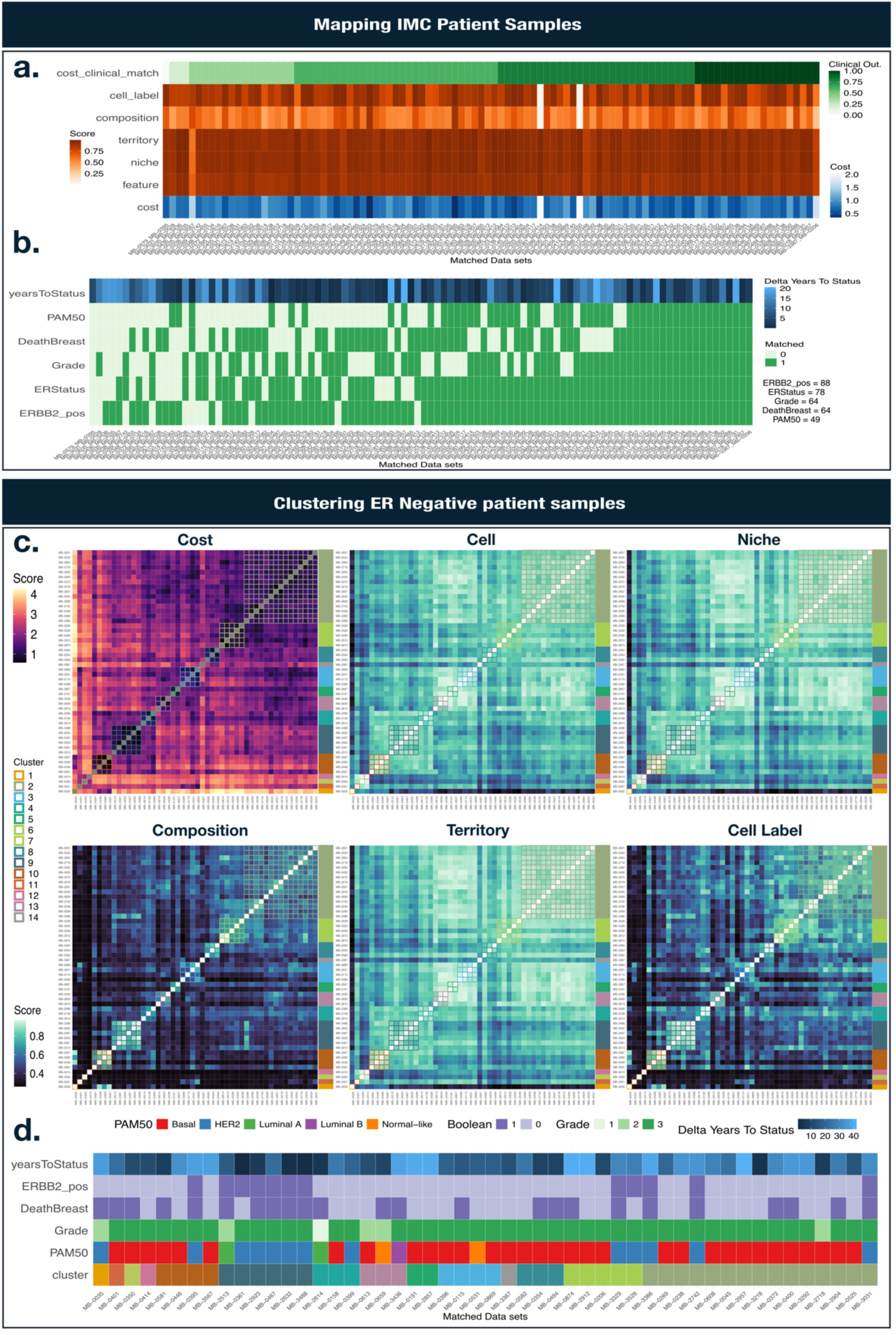
Mapping and clustering samples. (a) Scores of best matching sample pair across 100 sampled breast cancer data sets. We obtained higher mapping scores in metrics linked to the cell state continuum. The number of matches with the clinical property is counted along the best match. (b) ERBB2_pos is the metric that is the most accurately recovered (89%) while PAM50 (cancer sub-type) is the most poorly predicted (49%). (c) Clustering using ER negative breast cancer samples. The distinction of these samples is mainly driven by cell label and niche composition. (d) Clinical metrics associated to each cluster. We use the term Boolean to define True (1) or False (0).

We further performed hierarchical clustering for the ER negative patients (n = 50) using our total cost as a measure of distance. In total, we found 14 ER negative sub-population. Some clusters were driven by cell composition and cell type labelling (clusters 10, 9, and 4). We did not observe as strong of a difference in mapping scores related to cell similarity, niche similarity, and territory similarity in these clusters. We checked which clinical metrics were associated with these clusters (Figure 5d). Clusters 10 and 9 were characterized by Basal and HER2 cancer sub-types respectively. Our results suggest that cell type label for surrounding cells can dictate clinical metric. Clustering results of ER positive patients are shown in Supplementary Figure 4 and 5.

## Discussion

With the growing volume of spatial data, the necessity to align and compare it becomes increasingly crucial. It is inevitable that we reconcile spatial datasets that are dissimilar in their spatial structure and/or are collected using different technologies. Additionally, we need to compare spatial data obtained across biological processes or developmental time points. To address the challenge of mapping cell states across heterogenous samples, we developed a new tool in the framework of Vesalius. Vesalius differs from previous spatial matching tools that are designed to align data from adjacent tissues including PASTE^26^ and GPSA^28^. The many-to-many cell matching framework is similar to scRNA-seq mapping tools CytoSpace^30^ and Tangram^30^. However, inclusion of spatial and biological contexts such as cell, niche, and tissue territory similarities in Vesalius enabled effective mapping of spatial data across samples, technologies, resolutions, and time.

Our benchmarking results emphasize the algorithmic advantages against other approaches. In contrast to tools such as CytoSpace^30^ and Tangram^30^ which do not account for spatial context, Vesalius is able to recover the spatial distribution of cells in all regimes we tested, indicating that niche information is crucial in mapping spatial data. PASTE^26^ and GPSA^28^, developed for spatial alignment with an assumption of tissue structural similarity, are not suited for mapping cells across heterogenous samples. Even though SLAT^31^ was developed to map heterogeneous spatial data, it did not perform well in our test maybe it allows only limited degree of structural changes.

We showed that Vesalius successfully mapped sequence-based Slide-seqV2 and image-based seqFISH data. For each task, we used the context that can be obtained easily from the dataset. In Slide-seqV2, for instance, we used cell, niche, territories. In seqFISH, we used cell similarity, niche similarity, and niche composition. Vesalius integrates the mapping results by highlighting how different biological features participate in the spatial mapping. The mapping scores indicate that while many cells shared the same niche composition, the gene expression profiles of those niches were much more nuanced.

It is also worth noting that Vesalius performs cross-technology alignment. The alignment between image-based seqFISH and sequence-based Stereo-seq in mouse embryos showed remarkable matches in the corresponding cell types (Figure 3e). Mapping VisiumHD data onto low resolution Visium assay emphasizes the usefulness of Vesalius in integrating the existing public dataset (Figure 3e).

With its mapping, Vesalius can be utilized in various ways. First, Vesalius showed a remarkable ability to map cells across time and tissue states by recovering expected tissue structure during Axolotl brain regeneration and during murine embryonic development. Our integrated territory approach allowed us to underline genes being activated at the zone of injury. Brain cells in the mouse embryo demonstrated spatial sub-populations which lead to the discovery of spatial-temporal decoupling events related to stem cell differentiation. Second, we demonstrate that mapping cost constructed from a variety of biological features can be used to cluster samples and discover patient sub-populations. Our multi-context and multi-scale cost matrix provides a convenient metric to compare samples in terms of overall spatial similarities in IMC breast cancer data. The best matching sample pairs showed a remarkable concordance with clinical metrics such as ER status and ERRB2 positive patients even though we used only 37 protein markers. These results suggest that patient status can be at least partially encoded in a cancer’s multi-scale organization.

Vesalius is a versatile tool designed to map cell states across diverse samples at various scales. By incorporating spatial information, Vesalius offers a deeper understanding of cellular behavior in different contexts. We highlight the significance of spatial context and cell states in studying brain regeneration, development, and patient classification. As more data becomes available, the value of spatial mapping will continue to grow.

## Methods

### Mapping cells across samples

To map cells across samples, Vesalius employs the Jonker-Volgenant algorithm^32^, a variant of the LAP. This algorithm requires less computing cost (*O*(*n*^3^)) than the Hungarian Solver algorithm (Kuhn-Munkres algorithm – *O*(*n*^4^)). The algorithm is to map “workers” to “task” while minimizing the overall cost. To allow many-to-many matching of cells, we implemented the solver in a divide-and-conquer framework which also provides a speed up for large batches through parallelization.

Based on the requested batch size, we randomly sample cells from the reference and the query. Once a cell has been selected, it is removed from the sampling pool to ensure that all cells will be selected at least once. In the case where the number of cells in one data set is smaller than in the other, we add a cell padding where a new set of duplicated cells will be concatenated to the original selection. The padding will be added to the smaller data set and will be made to match the batch size or the number of cells in the other data set whichever is smaller. The LAP solver will be applied across each batch and the final mapping results will be concatenated together. If a one-to-one matching is required, duplicates matching events can be removed only retaining matching pairs with the lowest overall cost.

Since the matching will depend on the cells selected during sampling, the batching and solving steps can be applied across multiple epochs. At each epoch, new random batches will be selected and mapped. Then, the new cells pairs are compared to previous cell pairs and will be retained if the matching results in a lower matching cost. The final output is a many-to-many mapping of cells across samples alongside mapping cost, epoch, and mapping scores of each cell pair.

### Computing cost

To solve the LAP, we compute a total cost matrix constructed from comparing cells through a variety of biological features. The default features used by Vesalius are the following:

- *Cell similarity*: compares the gene expression of cells between each sample.
- *Niche similarity*: compares the gene expression of cell niches between each sample.

Optionally:

- *Territory similarity*: compares the gene expression of territories between each sample.
- *Cell label*: checks whether both cells share the same cell type label. If the labels are shared the pair will receive a score of 1 otherwise 0.
- *Composition similarity*: compares the cell type composition of cell niches between each sample.
- *Custom similarity*: compares cells between samples using user defined similarities.

The total cost matrix is built through pair-wise summation of the reciprocal of each similarity matrix provided. Let *A*, and *B* be our cell similarity and niche similarity scores matrices and *C* be our final cost matrix, then:

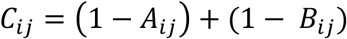

where *i* and *j* are the query cell *i* and reference cell *j*. The same process is applied to all matrices that are requested or provided.

In the case of continuous features (cell, niche, and territory), we first extract a gene expression signal. The signal represents a gene expression profile (normalized) of all genes, highly variable genes, or a custom gene set. Alternatively, embeddings values (PCA, UMAP, LSI, NMF) can also be used to define the gene expression signal. By default, and across all analysis presented here, we used the highly variable genes. Since genes might not perfectly overlap between data sets, we select the genes which lie at the intersection of gene expression signals for each data set. Next, we compute a Pearson’s correlation coefficient between expression signals of each potential cell pair:

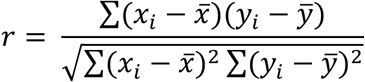

where *x_i_* and *y_i_* are the expression of genes *i* in sample *x* and *y* respectively. *x̅* and *y*6 are the mean expression of cell *x* and cell *y*.

For categorical features such as niche composition, we compute a frequency aware Jaccard index:

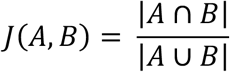

We assign a unique label to duplicated cell type labels. If cell type A is present more than once, the first instance the label will be A, the second instance A.1, and so on.

The scores are placed in a similarity score matrix where rows are the query cells/spots and columns are the reference cells/spots. This layout conventions can be used for any custom matrix that is provided by the user.

### Extracting Niches

For each cell, we define a niche as the spatially neighboring cells using the following methods:

- *KNN*: K-Nearest Neighbors
- *Graph*: using Voronoi Tessellation, we define a neighborhood graph and select niches by considering graph depth from a center cell. e.g. direct neighbors will have a graph depth of 1.
- *Radius*: All cells within a certain radius of a center cell are defined as this cell’s niche.

Niche expression is then defined as the averaged expression of all cells present in that niche. The composition scores (cell type composition) use the same niches.

### Extracting Territories

Originally, Vesalius was designed to detect tissue territories from spatial transcriptomics data using image processing techniques. We have since improved the territory detection abilities. In addition to a wider range of dimensionality reduction techniques (PCA, UMAP, Latent Semantic Indexing (LSI), Non-Negative Matrix Factorization (NMF), custom) and normalization methods (log normalization, SCtransform, Term-Frequency Inverse Document Frequency (TF-IDF), custom), we have adapted the image processing steps of Vesalius to function on image tensors instead of RGB formatted images. This upgrade allows us to apply the image processing steps to all latent spaces dimensions simultaneously and in turn improve our ability to recover tissue territories.

In summary, spatial transcriptomics data is converted into a grey scale image stack. We normalized counts and extract highly variable features. With these features, we compute a reduced dimension latent space for each cell. In parallel, we expand punctual coordinates using Voronoi tessellation and rasterize the resulting tiles to produce a grey scale image for each latent space dimension. This image stack can be processed using a variety of image processing techniques such as histogram equalization, image smoothing, and image segmentation. Color segments can be further divided to distinguish spatially distinct color segments. The details of this process are discussed in our previous publication^22^.

The territories isolated by Vesalius are used to compare the expression of cells across larger scale spatial domains. We compare the average expression of the territory in which a cell finds itself across samples.

### Spatial Simulations and benchmarking

To benchmark our mapping approach, we elected to use simulated data since they provide a strong ground truth against which we can compare performance. We used 3 regimes: *circle*, *layer* and*, dropped*.

For the *circle* regime, we created 5 circular territories of random size placed at random within a background territory. We allowed overlaps between territories but ensured that all territories were present across 12 samples. The circle regime consists of 5000 cells and 2000 simulated genes with a p = 0.3 probability of gene expression differences between cell populations in each sample. Each circle (and the background territory) contains 2 different “cell types” for a total of 12 cell populations. We used the splatter^39^ package to generate synthetic single-cell-like count data for all samples. All cells for each cell type across each sample are taken from the same synthetic distribution.

For the *layer* regime, we employed a similar data generation strategy for the 12 samples with a few key differences. First, only a single circular territory is placed at random in the background territory. This territory is then subdivided into 5 layers with each layer containing 2 cell types. The probability of gene expression difference between cell types in the layers is reduced to 0.05. The *dropped* regime follows a similar pattern as the layered regime, but we generate 2 circular territories divided into 3 layers. Each layer can contain a varying number of cells in varying proportions. While cells will be taken from the same statistical distribution for each cell types, in the *dropped* regime, not all cells will be shared between all data sets.

To facilitate reproducibility and replicability, we wrapped our simulation approach in an R package - oneiric - available of GitHub. The package vignette will generate the data sets used for benchmarking as well as a demonstration on how to create other regimes.

We contrast our mapping approach to single cell to spatial mapping tools and spatial alignment tools. More specifically, we compared Vesalius to CytoSpace^32^, Tangram^32^, PASTE^26^, GPSA^28^, and SLAT^31^. It is of note that CytoSpace requires perfect matching of cell type labels between reference and query and only finds an optimal mapping of cells between the same cell type labels. To demonstrate how this strategy differs from our context specific approach to cell mapping, we provided CytoSpace with a single cell type label for all cells.

To quantitively assess the performance of cell mapping between samples, we computed an adjusted rand index^40^ (ARI) and variation of information^41^ (VI) between the cell labels of the query and the reference. Since PASTE and GPSA do not match cells, we first computed the nearest cell neighbor between reference and query and compared their cell type labels. While this approach disadvantages PASTE and GPSA, our objective is to demonstrate that context mapping is a fundamentally different task than spatial alignment.

### Cross sample mapping

We tested mapping cells across samples from the same technologies using Slide-seq V2^8^ and seqFISH^33^. Prior to mapping, we filtered the Slide-seq sections from mouse hippocampus to retain cells that contained at least 200 counts and genes with at least 100 counts. We pre-processed each slide independently using Vesalius. First, counts are log normalized; then we generated grey scale color embeddings from PCA (30 PCs) using the top 2000 variable features. We processed the image stack by equalizing the color histogram and smoothing each image. We then isolated tissue territories through image segmentation. We then mapped the query (Puck_190921_21) onto the reference (Puck_200115_08) using cell similarity, niche similarity, and territory similarity across samples. We defined the niche using K-NN with k = 30. We set the batch size to 10000 and optimized the mapping across 10 epochs. We used variable features to define cell signal. Slide-seq analysis code is provide in a dedicated GitHub repository (see Code Availability).

In the case of seqFISH mouse embryo data, we filtered out all low-quality cells as described by the authors. We elected to map “embryo1” to “embryo3” both representing different mouse embryo sections. We applied a similar pipeline as Slide-Seq V2 but with parameters adapted to the data. We defined niche using K-NN with k= 6. We set the batch size to 5000 and compared cells and niches using all gene features since this would be equivalent to using variable features (351 total genes). In addition, we demonstrated the effect of varying cost matrices by running the same analysis pipeline but with different similarity matrix combinations. Since the published seqFISH data contains cell type labels, we used these labels to quantitatively assess the mapping performance through metrics such as Adjusted Rand Index and Variation of Information. seqFISH analysis code is provide in a dedicated GitHub repository (see Code Availability).

### Cross technology and cross resolution mapping

To compare cells across technologies, we processed seqFISH^33^ and Stereo-seq data^9^. seqFISH data was filtered by removing low-quality cells as described by the authors. Next, we pre-processed both data sets using Vesalius by log-normalizing the counts and producing a reduced dimension latent space using PCA (PCs = 30). Our cross-technology comparison only used cell similarity and niche similarity as such no further processing was required. We mapped cells from Stereo-seq onto seqFISH using graph to define neighborhood with a depth of 2. We optimized the cell matching in batches of 10 000 cells across 10 epochs.

Matching cells across resolutions required extra steps for data preparation. First, we downsampled the Visium HD mouse brain data to retain 200 000 cells. We then used RCTD in doublet mode to deconvolve and assign cell type labels. As a reference, we used the labelled scRNA data provided by^42^. We remove all cells that contained the “reject” tag. For all remaining cells, we annotated cells with one or two labels depending on how RCTD defined them (singlet or doublet). After this filtering, the VisiumHD data contained 84 287 cells. We also used RCTD to deconvolve the Visium mouse brain data with the same reference data set^42^. In this instance, we only retained cell type labels that were considered as “high confidence”.

For cross-resolution mapping, we constructed the cost matrix using niche and territory. We defined territories by processing each data set through the Vesalius pipeline which includes color histogram equalization, image smoothing, and image segmentation. The parameter details are available in the analysis scripts deposited in our dedicated GitHub page (see Code Availability). We defined niche using a radius from the center cell where the radius is defined as half of the scaled distance between Visium Spots. Cell matching was achieved across 20 epochs with a batch size equal to the number of Visium spots (n = 2310). We used log normalized variable features as the cell expression signal to compare. Once the matching was completed, we ran the mapped data set through the Vesalius pipeline to retrieve spatial domains, once with a jitter added to the coordinates and once without any jitter. The jitter enables each mapped cell to have a distinct spatial coordinate while no jitter will merge the mapped cells into Visium-like spots.

To compare the cell types mapped across resolution, we computed a Jaccard index between cell types present in the Visium Spot and the VisiumHD spots mapped to Visium. The higher the index the higher the concordance in cell types being mapped.

### Count Integration

Once cells have been matched across samples, we integrated the counts and developed an integrated tissue analysis framework. To integrate counts, we used the Seurat implementation of Canonical Correlation Analysis^43^ (CCA) which returns scaled and integrated counts and a common latent space between samples. The integrated latent space is directly used to generate Vesalius images upon which our pipeline can be applied. To avoid duplicated coordinates, we add a jitter to the duplicated coordinates only. The resulting territories are the territories emerging from this joint latent space which can be used for spatially resolved differentially gene expression analysis.

For inter sample differential gene expression analysis, Vesalius can compare the cells from the integrated count matrix using either territories or cells as a grouping criterion. By default, we used Wilcoxon ranked sum test with the p-value threshold set at 0.05 after correcting for multiple testing (FDR).

### Cell mapping clusters

We developed a cell clustering method which clusters query cells based on which reference cells they tend to co-cluster to. Our default clustering approach is based on hierarchal clustering with community-based clustering approaches (Leiden and Louvain) also available. First, we define the metric matrix that will be used for clustering. This approach follows the same pair-wise summation of similarity scores used during cost matrix creation. By default, we use the cell similarity only.

Next, for each cell, we compute the order of preferential mapping (*i.e* lowest mapping cost) and convert the numeric value into a categorical string. We select the top *n* (default set at n = 30) matches and use this categorical vector as input to compute a Jaccard Index. The higher the score the more likely 2 query cells will map the same set of reference cells. The reciprocal of the Jaccard Index is used as the distance matrix during hierarchical clustering. For clarity, we used a fixed number of clusters with k = 5.

### Spatial proteomics *in situ* Mass Cytometry sample preparation and processing

To demonstrate the utility of Vesalius, we mapped spatial *in situ* Mass Cytometry (IMC) data taken from human breast cancer patient samples^38^. Out of the 709 samples, we filtered the samples to ensure that they contained at least 1000 cells. We also filtered out samples that contained NA’s in the clinical metrics provided by the authors (ERStatus, ERBB2_pos, grade, deathBreast, PAM50, and YearsToStatus). From the remaining samples, we randomly sampled 50 ER positive and 50 ER negative samples.

Next, we extracted the expression values for the 37 markers measured that we formatted into a count matrix for Vesalius. We processed every sample using the Vesalius pipeline including log-normalization, dimensionality reduction (PCA) and images processing, and territory isolation. The details of the parameters used are contained in our analysis scripts deposited in a dedicated GitHub page (see Code Availability).

We mapped cells between samples using a cost matrix built from cell similarity, niche similarity, territory similarity, cell label, and niche composition. We optimized the matching pairs across 10 epochs with a batch size equal to smallest data set minus one (or 1000 - 1 cells if both data sets were larger than a 1000 cells). The neighborhood was defined through our graph method with a depth of 2. For each mapping event, we extracted the mean cost and mean mapping scores.

For each sample, we selected its best matching pair (excluding self-mapping) by taking the pair with the lowest overall cost. From this best matching pair, we computed the overlap between the clinical metrics to check whether our mapping strategy could recover patient level information. More specifically, we checked if we obtained the same label between the reference and query. In the case of Grade and PAM50, there are multiple labels: “1”, ”2”, “3” for Grade and “Basal”,”HER2”, “Luminal A”, “Luminal B”, “Normal-like” for PAM50. The delta *YearsToDeath* was calculated by taking the absolute value of the difference in years between query and reference.

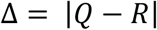

### Sample mapping clusters

To cluster samples, we first mapped samples to each other, extracted their average cost, and averaged mapping scores across all cells in those samples. Since we are mapping sample to each other, we can directly use the mapping cost matrix as input to the hierarchal clustering algorithm. In this case, we used height to define the number of cluster cutoff (h = 0.5). Once we defined clusters for each sample, we grouped them for visualization.

## Data Availability

*Simulated data*: https://github.com/WonLab-CS/oneiric

*Slide-seq V2*: https://singlecell.broadinstitute.org/single_cell/study/SCP815/sensitive-spatial-genome-wide-expression-profiling-at-cellular-resolution#study-summary

*seqFISH*: https://content.cruk.cam.ac.uk/jmlab/SpatialMouseAtlas2020/

*MOSTA*: https://db.cngb.org/stomics/mosta/stereo.seq/

*Visium* : https://cf.10xgenomics.com/samples/spatial-exp/2.0.0/CytAssist_FFPE_Mouse_Brain_Rep1/CytAssist_FFPE_Mouse_Brain_Rep1_web_summary.html

*VisiumHD* : https://www.10xgenomics.com/datasets/visium-hd-cytassist-gene-expression-libraries-of-mouse-brain-he

scRNA: Molecular Diversity and Specializations among the Cells of the Adult Mouse Brain.^42^

*ARTISTA* : https://db.cngb.org/stomics/artista/

*IMC*: Breast tumor microenvironment structures are associated with genomic features and clinical outcome^38^

## Code Availability

Vesalius 2.0 is available on GitHub: https://github.com/WonLab-CS/Vesalius

Oneiric is available on GitHub: https://github.com/WonLab-CS/oneiric

All analysis code is available on GitHib: https://github.com/WonLab-CS/Vesalius_analysis

## Supplementary Material

Supplementary plots are contained in the Supplementary manuscript.

List of differentially expressed genes for Axolotl brain regeneration and murine brain development are contained within Supplementary Table-1 and Supplementary Table-2. (csv files)

## Acknowledgments

The authors gratefully acknowledge the support of Cedars-Sinai Medical Center through institutional funding.

## Supplementary figure

**Figure S1:**
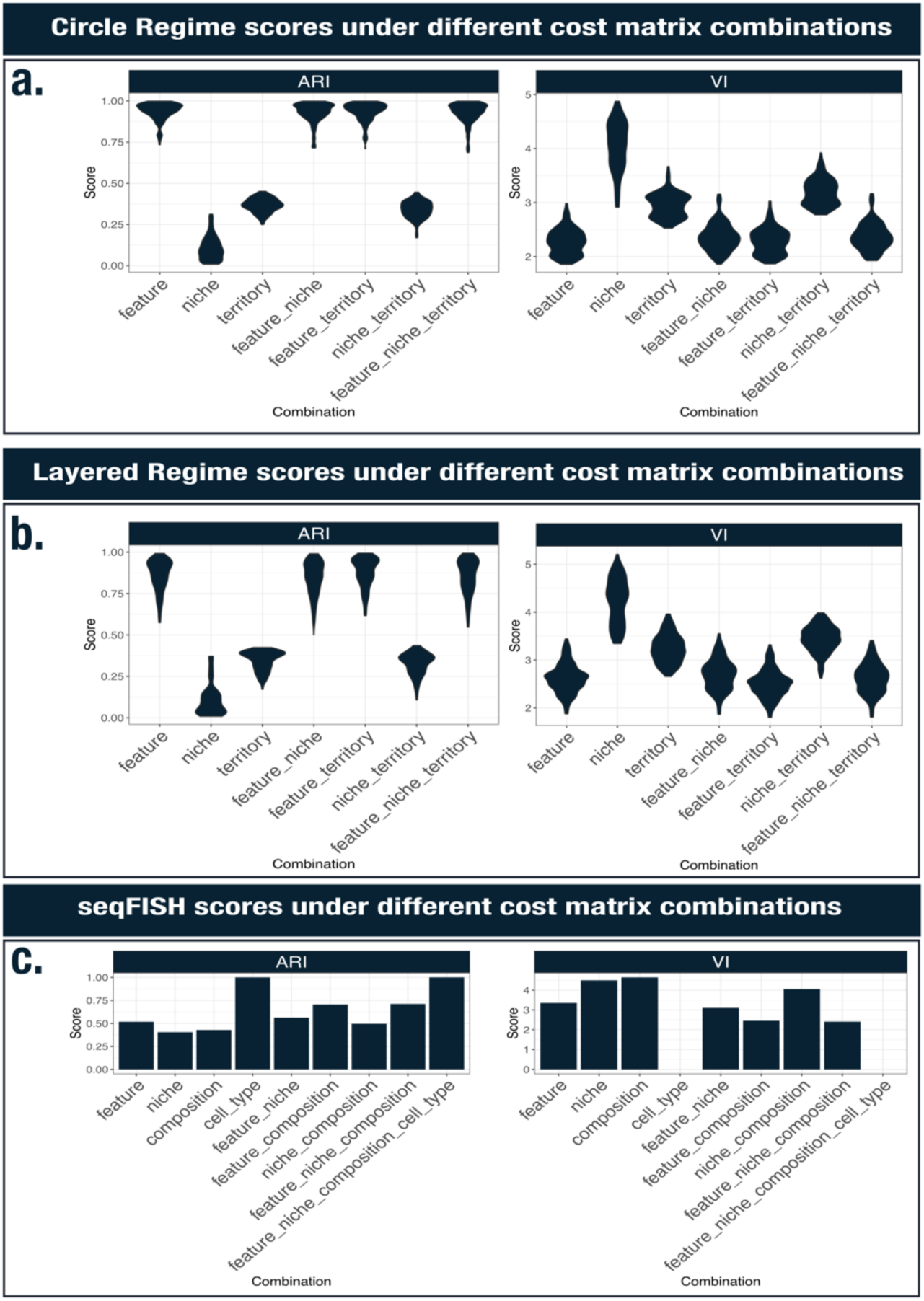
Mapping performance under different cost matrix selection. (a) Mapping performance using an Adjusted Rand Index (ARI) and Variation of Information (VI) using a variety of cost matrix combination in the synthetic circle regime. (b) Mapping performance using an Adjusted Rand Index (ARI) and Variation of Information (VI) using a variety of cost matrix combination in the synthetic layered regime. (c) Mapping performance using an Adjusted Rand Index (ARI) and Variation of Information (VI) using a variety of cost matrix combination in the seqFISH mouse embryo data.

**Figure S2:**
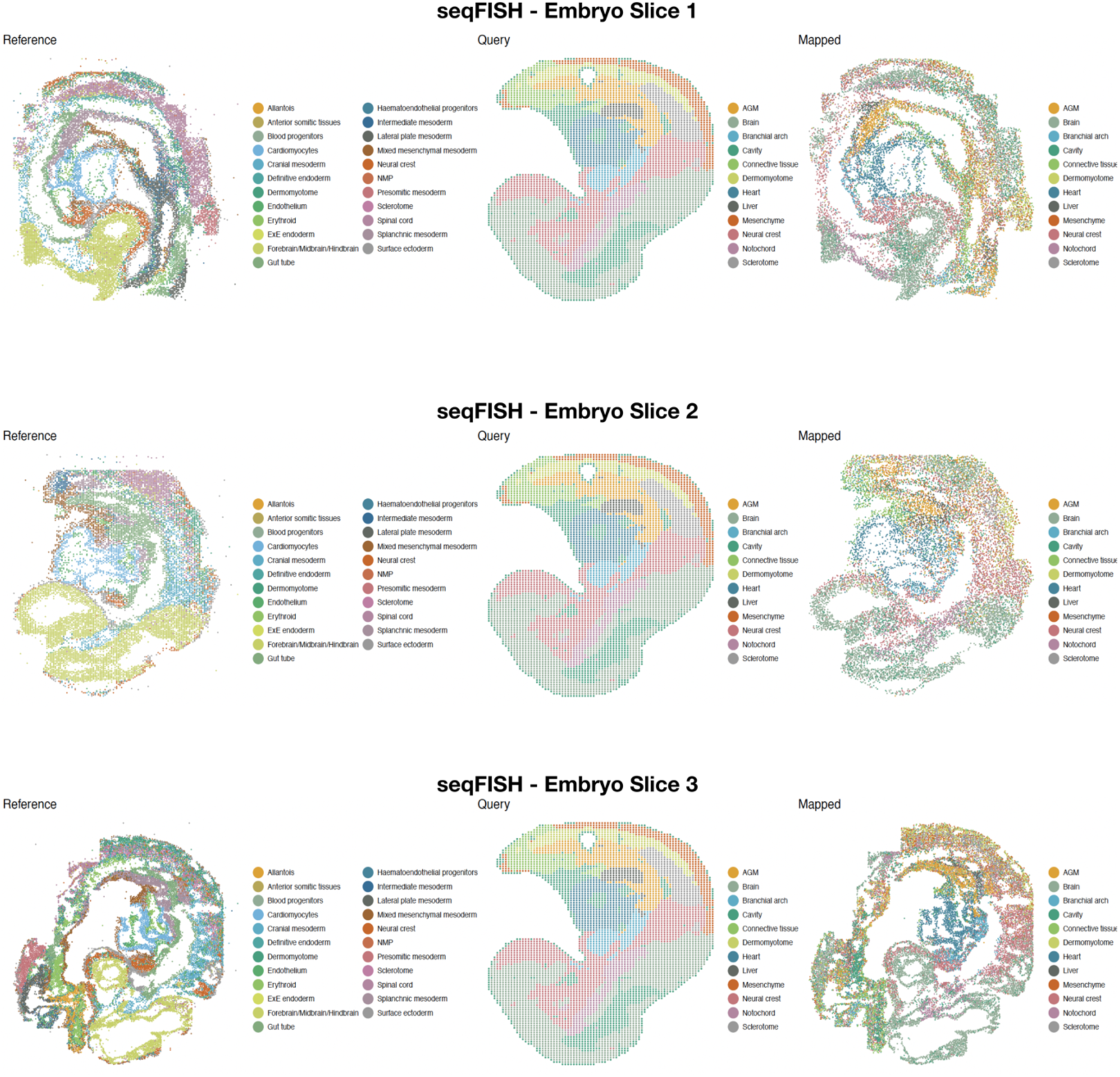
Mapping of Stereo-seq onto all 3 seqFISH Mouse embryo sections.

**Figure S3:**
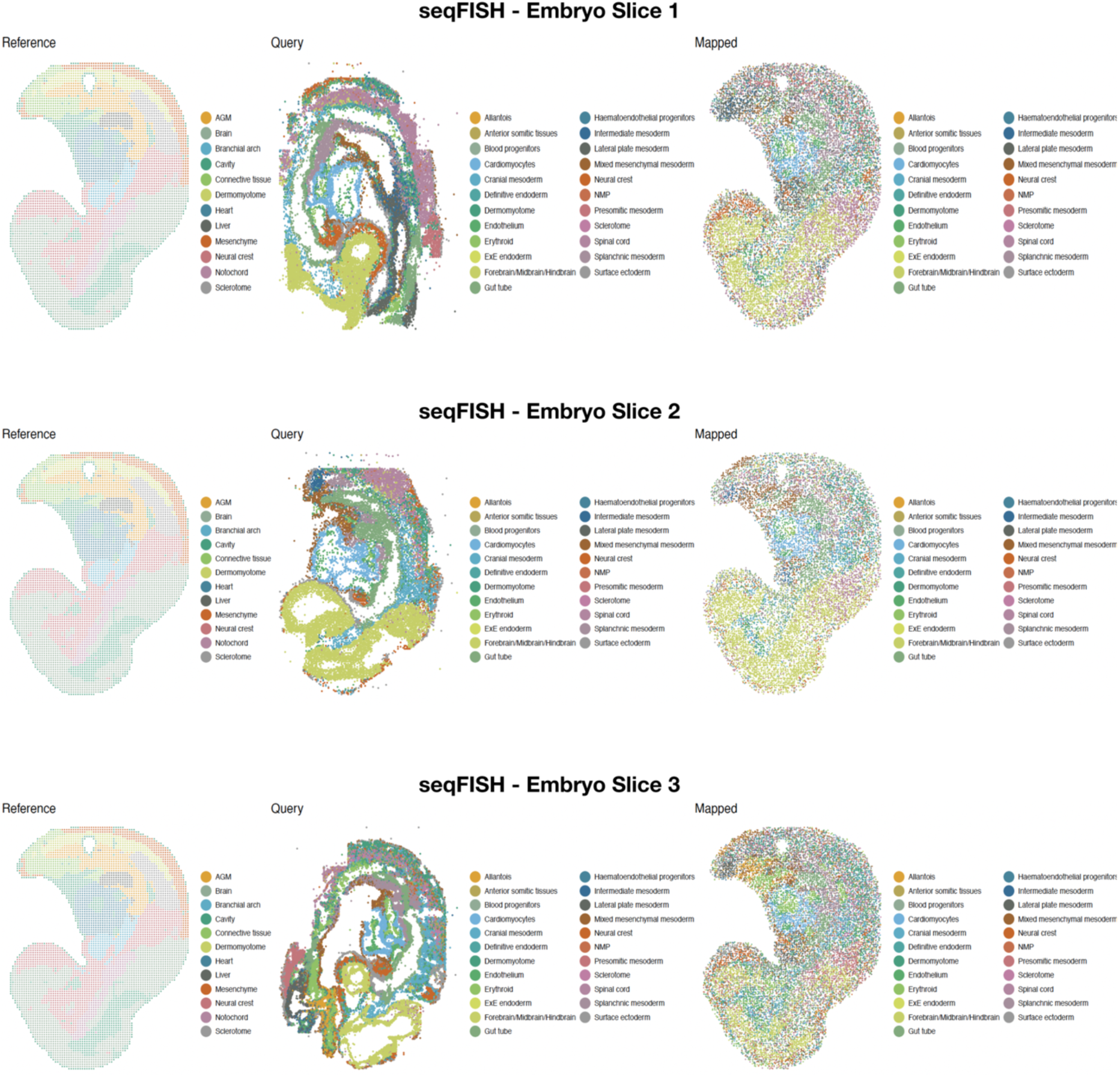
Mapping of seqFISH data onto Stereo-seq data.

**Figure S4:**
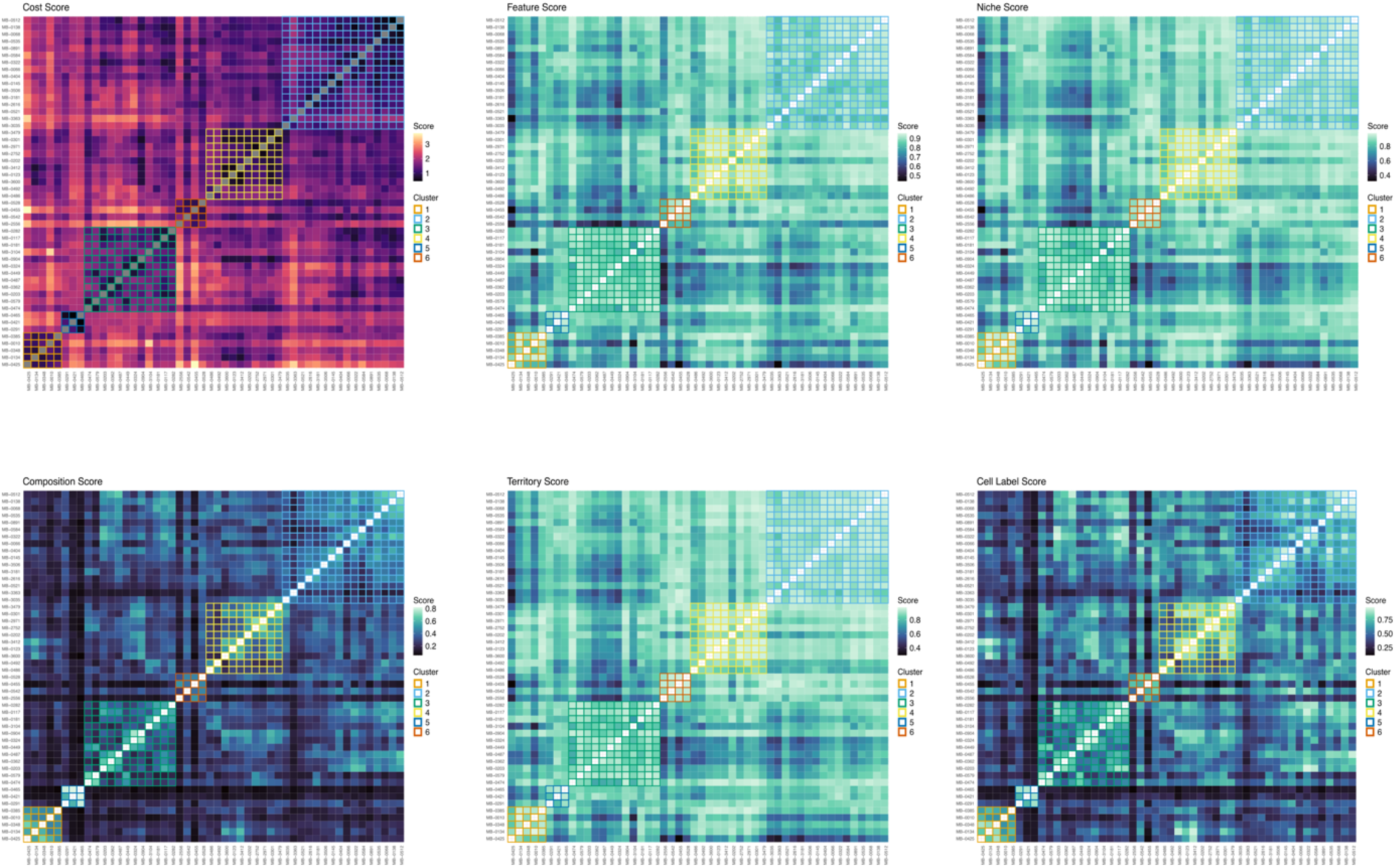
ER positive patient clustering.

**Figure S5:**
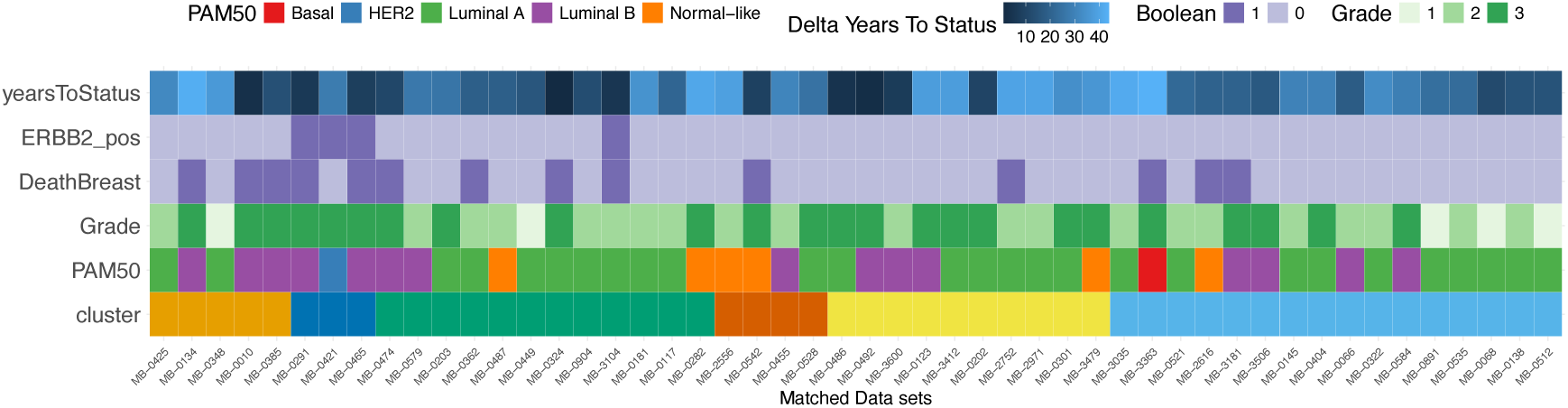
Clinical metrics associated with the clustering of ER positive patients. We use Boolean to represent True (1) and False (0) values.

